# The spatiotemporal evolution of TMS-evoked potentials reflects direct cortical activation

**DOI:** 10.1101/2025.06.25.661535

**Authors:** Matteo Fecchio, Simone Russo, Bruno Andry Couto, Ezequiel Mikulan, Andrea Pigorini, Giulia Furregoni, Gabriel Hassan, Sasha D’Ambrosio, Michela Solbiati, Alessandro Viganò, Sara Parmigiani, Simone Sarasso, Silvia Casarotto, Marcello Massimini, Adenauer Girardi Casali, Mario Rosanova

## Abstract

Transcranial Magnetic Stimulation (TMS) evokes electroencephalographic (EEG) responses that can persist for hundreds of milliseconds. While the first 80 ms after the pulse are widely accepted to reflect genuine cortical responses to TMS, later components have mainly been attributed to the effects of sensory co-stimulations. Here we reappraise this view by investigating the target-specificity of the spatiotemporal evolution of TMS-evoked potentials (TEPs). To this end, we compared TEPs elicited by targeting the premotor and primary motor cortices in 16 healthy subjects, under conditions designed to optimize TMS effectiveness on the cortex while minimizing peripheral confounds. As a counterfactual, we conducted the same comparison on the EEG responses evoked by realistic sham TMS and high-intensity somatosensory scalp stimulation. We found that EEG responses to motor and premotor TMS can exhibit distinct spatiotemporal evolutions, lasting up to 300 ms, both at the group and single-subject levels. These differences were absent or marginally detectable in both realistic sham TMS and high-intensity somatosensory scalp stimulation. Our findings suggest that, when effectiveness is optimized and peripheral confounds are controlled, TMS elicits specific long-lasting genuine EEG responses that reflect the initial engagement of specific cortical targets. These results challenge previous assumptions and highlight how TMS-EEG can be reliably used to assess large-scale properties within corticothalamic networks.

## INTRODUCTION

Transcranial Magnetic Stimulation combined with Electroencephalogram (TMS-EEG) allows probing cortical reactivity and connectivity directly and non-invasively, offering high potential for research and clinical applications [1–3]. However, its clinical adoption has been hindered by the lack of standardized procedures for collecting and analyzing TMS-evoked potentials (TEPs) [4–6].

For example, a variety of methods are used to set stimulation parameters and data acquisition criteria. Different research groups calibrate TMS intensity in various ways, either based on motor output, phosphenes perception, induced electric field strength, or EEG response amplitude, leading to inconsistent results [5,6]. Furthermore, biological confounds, such as TMS-related cranial muscle activations and auditory evoked potentials (AEP), are addressed either using offline signal processing methods [7–10] or applying preventive procedures during recording [11–14]. Finally, although several sham TMS protocols exist to account for auditory and somatosensory effects on TEPs [15], no standard approach has been agreed upon yet.

As a result, the TMS-EEG community is still debating the extent to which TEPs reflect genuine responses to direct cortical stimulation, unconfounded by sensory co-stimulations [16,17]. For instance, Conde and colleagues [18] found that EEG responses to TMS targeting different cortical areas share common spatiotemporal features with corresponding realistic sham stimulations as early as 20 ms post-stimulus, indicating contamination by the effects of peripheral co-stimulation. Following studies suggest that genuine TEPs can last until 80 ms after the TMS [19–26], while others extend this limit to later latencies [27–32].

Here, we address this open issue by comparing TEPs collected by targeting nearby cortical areas (i.e., the premotor -PM- and primary motor -M1- areas). We assume that the spatiotemporal evolution of the two EEG responses reflects sensory effects. Therefore we consider the null hypothesis that it should not differ, given that shifting skin and auditory stimulation across short scalp distances is not expected to produce significant changes in sensory representations [33]. First, we challenge this hypothesis, under experimental conditions designed to optimize cortical activation by TMS (effective TMS, eTMS) while minimizing peripheral sensory co-stimulation [11,34]. As a counterfactual, we compared EEG responses to stimulation of the same targets in the absence of direct cortical engagement, using two types of sensory stimulation: a realistic sham (RS-TMS), combining electric scalp and auditory stimulations to replicate TMS-related sensory co-stimulations; and a high-intensity electrical scalp stimulation (HI-ES), delivering the highest tolerable scalp stimulation below pain threshold.

We found that EEG responses evoked by eTMS of adjacent cortical targets show significant spatiotemporal differences lasting up to 300 ms, thus rejecting the null hypothesis. In contrast, EEG responses to sensory stimulation showed no or only marginal differences across the same targets. Taken together, these findings suggest that TEPs can reflect specific cortical activations even at later latencies, with little to no contribution from peripheral confounds.

## MATERIALS AND METHODS

### Study participants

We enrolled sixteen healthy subjects (n=16; 4 female, 3 left-handed, age: 25-51 years, Table S1). Exclusion criteria included neurological or psychiatric disorders, current CNS-active drugs use, substance abuse, and contraindications to TMS. The Ethics Committee Milano Area A approved the study, and all subjects provided written informed consent.

### Experimental design and protocol

We systematically compared the EEG responses between PM and M1 stimulation targets, at both group and single-subject level, and for each stimulation type: eTMS, RS-TMS, and HI-ES (Figure 1).

**Figure 1.**
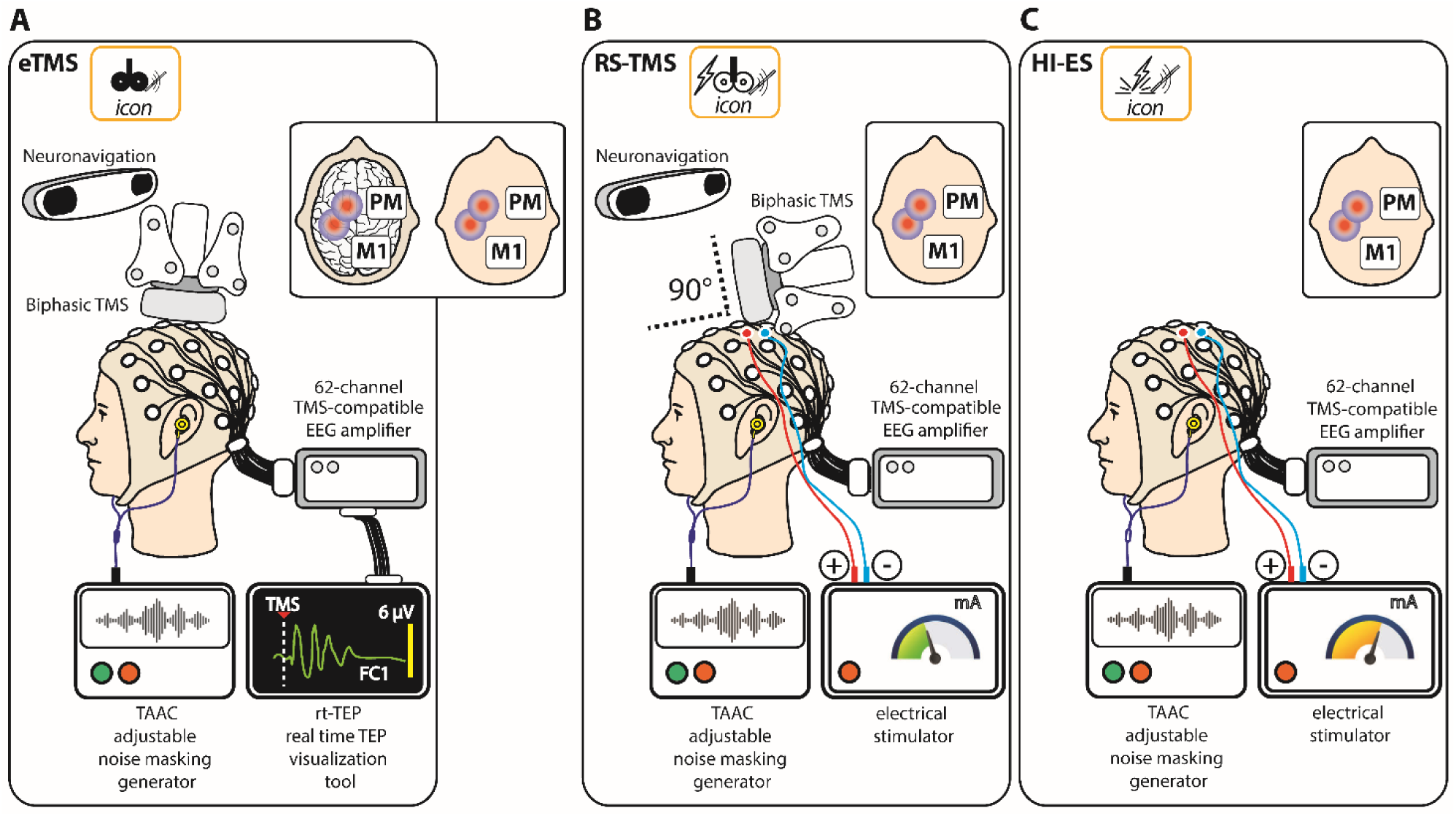
Schematic representation of the experimental setup for effective TMS, realistic sham TMS, and high-intensity electrical stimulation. In each panel, the acronym and the icon (within the orange frame) for each stimulation condition are reported in the top-left corner. EEG was acquired with a 62-electrode cap connected to a TMS-compatible amplifier. The insets show schematic representations of the target locations for PM and M1. Noise masking was played in all conditions (bottom left of each panel). **(A)** Biphasic TMS pulses were delivered over the left PM and M1 cortical areas using a neuronavigated figure-of-eight TMS coil. Rt-TEP, a real-time visualization tool (bottom right) was used to optimize TMS parameters and evoke early TEP components (< 50 ms) with amplitudes ≥ 6 µV beneath the coil. **(B)** RS-TMS was designed to mimic scalp and auditory sensations associated with TMS without actual cortical stimulation. Electrical stimulation was applied to scalp targets overlying the same cortical areas targeted with TMS. Pairs of stimulating electrodes (blue and red), spaced 2.5 cm apart [18], were positioned to match the TMS coil location and the direction of the TMS-induced electric field. The TMS coil, rotated 90° in the sagittal plane, was kept in contact with the scalp over the target area, a position known to effectively eliciting TMS-related AEPs and preserving coil vibrations [35]. TMS and electrical stimuli were delivered simultaneously, with intensity adjusted to match the scalp sensation of TMS based on participant’s reports (Supplementary Methods 4, Table S3). **(C)** High intensity electrical stimulation was applied to the same scalp targets used for RS-TMS, but without the TMS coil. Stimulus intensity was gradually increased to the highest level tolerated by each participant without inducing pain (Supplementary Methods 4, Table S3).

TEPs and EEG responses to RS-TMS and HI-ES were recorded in two separate sessions held on different days, both starting at 10 AM to minimize circadian effects. Each stimulation type required tailored experimental procedures (Supplementary Methods 1-5). Noise masking, whose parameters were adjusted on each subject’s TMS “click” perception [34] (Supplementary Methods 2), was played during all EEG recordings, regardless the stimulation condition. To collect TEPs, sensory confounders, such as AEP, scalp muscle contractions, and motor evoked potentials (MEPs) [35], were minimized (Supplementary Methods 3 and 5).

Participants were seated comfortably during recordings and rated discomfort and pain on a Numerical Rating Scale (NRS; Supplementary Methods 6, Figure S1, and Table S2) at the end of each measurement. At least 200 TMS pulses were delivered per each eTMS session, and 300 stimuli per each RS-TMS and HI-ES session. For all stimulation types, stimuli were randomly delivered at an interval jittering between 2000 and 2300 ms (0.5–0.4 Hz).

### EEG recording and processing

EEG signals were recorded in direct current (DC) mode at 5 kHz using a 62-electrode cap connected to a TMS-compatible amplifier (BrainAmp DC, Brain Products GmbH). Electrodes were referenced to the forehead, with impedances kept below 5 kOhm. Preprocessing, including ICA for artifact removal, was performed as described in the Supplementary Methods 7 and 8, and Figure S2. To avoid bias in statistical comparisons, the first 135 good epochs (the minimum across all recordings) were used to compute evoked potentials. Grand-average evoked potentials were calculated by averaging single-trial responses across subjects for each target and experimental condition (eTMS, RS-TMS, and HI-ES).

### Statistical analyses

For each experimental condition (eTMS, RS-TMS, and HI-ES), we compared evoked potentials across different targets at both the group- (grand-average) and single-subject levels using a non-parametric permutation test with cluster-based correction for multiple comparisons (FieldTrip Toolbox [36]; Supplementary Methods 9). Significant differences between targets were considered only in channels and time samples showing a significant evoked response for at least one target, as determined by a surrogate data approach based on circularly shifting trial sequences [37] (Supplementary Methods 10).

In addition, we extracted reproducible evoked components across epochs and subjects using group-Task Related Component Analysis (gTRCA) [38], as described by Couto and colleagues [39]. For each condition and target, we retained the first (i.e., most reproducible) gTRCA component together with its eigenvalue, which quantifies the degree of reproducibility at the group level. Statistical significance of each eigenvalue was assessed using the subject-based shifting method, with 500 permutations (p < 0.05).

Stimulation intensities were compared between targets for each condition using the paired Wilcoxon signed-rank test. A linear mixed-effects model (LMM) was used to assess the effects of experimental condition and stimulation target on the amplitude of late evoked responses and NRS outcomes (discomfort and pain). Condition and target were fixed factors, with a random intercept for each participant to account for repeated measures.

## RESULTS

We recorded TEPs, and EEG responses to RS-TMS, and HI-ES while stimulating left PM and left M1 in 16 healthy subjects. Data recorded during RS-TMS and HI-ES in two subjects were discarded due to large decay artifacts that could not be removed without significantly corrupting the EEG response. After titrating noise masking (Supplementary Methods 2), all subjects reported no auditory perception of the TMS “click” as confirmed by the absence of TMS-related AEPs (Figure S3) in eTMS and RS-TMS conditions.

Stimulation intensities for eTMS, RS-TMS, and HI-ES are reported in Table S3 for each subject and target. Notably, eTMS intensity (expressed as the percentage of the maximal stimulator output), which affects coil vibration and hence tactile scalp stimulation, was not significantly different between targets (Wilcoxon signed rank test, W=44.5, z=-1.22; p = 0.22).

Subjectively, eTMS resulted in low discomfort (NRS 0-4), and no pain (NRS = 0 in all but one subject, which scored 1). Notably, 13 out of 16 subjects reported a discomfort difference of 1 or less between PM and M1, with no significant differences in NRS values for discomfort or pain between targets. Despite RS-TMS intensity being matched to the TMS-related sensory stimulation of the scalp, discomfort and pain levels were higher during RS-TMS (Figure S1 and Tables S4 and S5), with a maximum of 6 for discomfort and 3 for pain. Only three subjects reported discomfort and pain differences greater than 1 between targets. During RS-TMS stimulation, all subjects reported that the coil touching the scalp reduced discomfort, which otherwise was perceived as “too focal and pricking.” Finally, as expected, subjects reported significantly higher discomfort and pain during HI-ES, with maximum scores of 9 and 4, respectively (Figure S1 and Tables S4 and S5). Only two subjects reported differences greater than 1 across targets (Figure S1 and Tables S2).

### eTMS evokes site-specific EEG responses

To test the null hypothesis that the spatiotemporal evolution of EEG responses from PM and M1 stimulation do not differ due to sensory contributions, we first compared TEPs at the group level.

Figure 2A and 2B show the grand-average TEPs for PM and M1, respectively (TEPs collected in one representative subject are reported in Figure S4A). The scalp voltage distribution of the first TEP component is asymmetrical, with a more frontal distribution in PM than M1. The Global Mean Field Power (GMFP) calculated for significant evoked responses (Supplementary Methods 10) shows different time courses, with peaks at distinct latencies for the two targets, and diverging topographies. Specifically, late PM TEP peaked at 80, 112, and 213 ms, with slightly lateralized topographies involving frontal electrodes over the stimulated hemisphere, while M1 TEP peaked at 85, 128, 180, and 313 ms, mainly showing a clearly lateralized topography over the stimulated hemisphere. Cluster-based statistical tests confirmed these differences, identifying significant clusters in both amplitude and phase domains, mainly over the stimulated hemisphere, persisting beyond 300 ms (Figure 2C and 2D).

**Figure 2.**
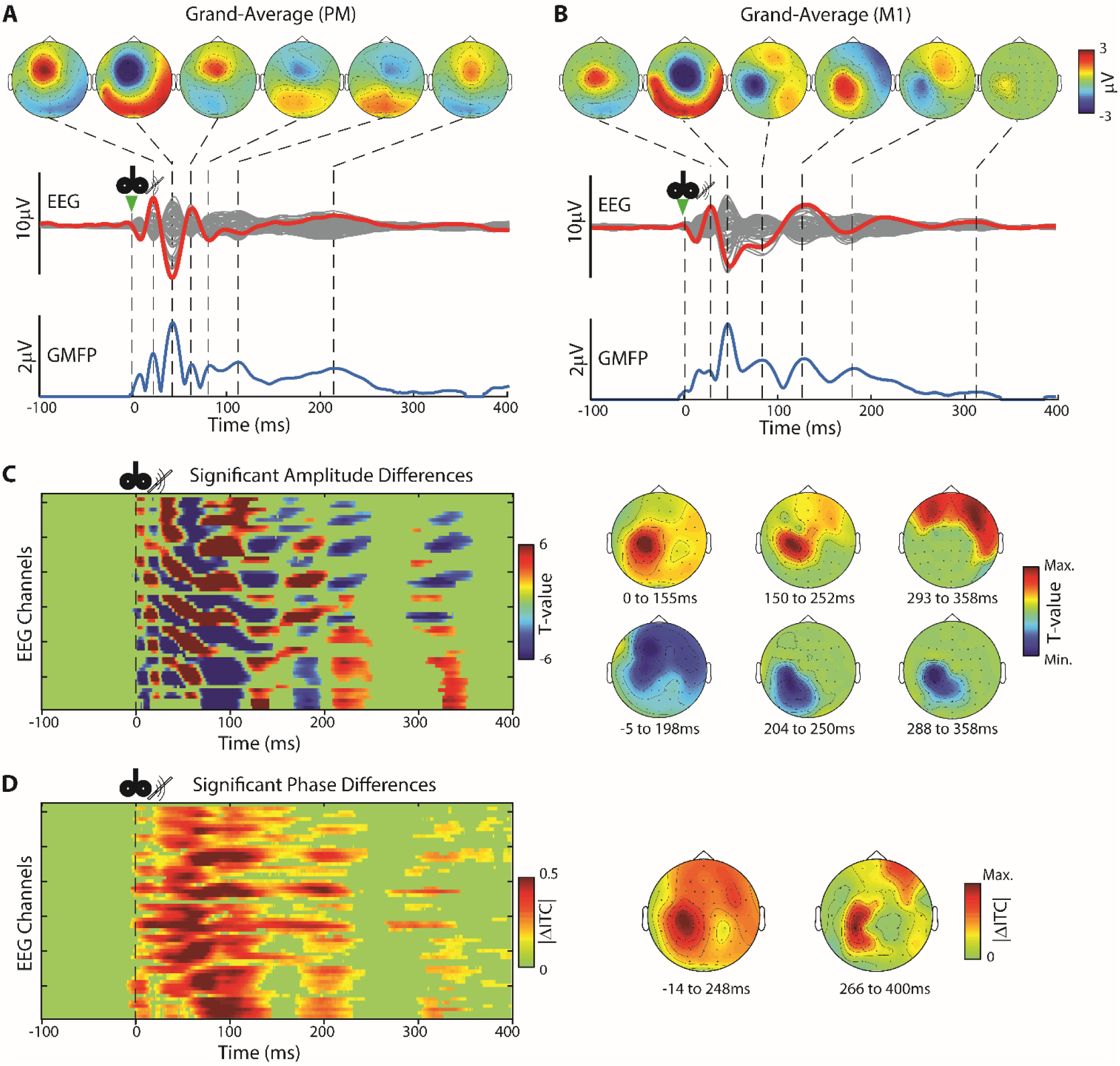
Grand average EEG responses to eTMS of PM and M1 differ significantly across multiple scalp regions and over long latency periods. **(A, B)** TEPs averaged across subjects following stimulation of the premotor (A) and motor (B) cortices are shown. The butterfly plot (middle row) shows all EEG channels superimposed (gray), with the red trace highlighting the EEG response of the channel closest to the stimulation site (FC1 for premotor and C1 for motor). GMFP values were calculated from significant TEPs and are displayed at the bottom row (blue). The topographical scalp voltage distributions (top row) are presented at the times of the GMFP peaks (dashed vertical lines). **(C, D)** Cluster-based statistical tests performed on the grand average TEPs reveal significant differences in amplitude (C) and phase (D) between responses evoked by premotor and motor stimulation. On the left, heatmaps showing the statistical results (T-values for amplitude and absolute difference in inter-trial coherence for phase) of significant clusters (blue and red for panel C, red for panel D) across EEG channels and times are displayed. Average topographical distributions of individual clusters are exhibited on the right, along with the corresponding time intervals. Six clusters are identified from amplitude comparisons, extending up to 358 ms, and two clusters from phase comparisons, extending up to 400 ms post-stimulation.

Additionally, to confirm that the differences between targets are consistent at the individual level, we conducted two complementary analyses. Figure 3 shows the waveforms and average scalp maps of the first (most reproducible) gTRCA component for PM and M1 TEPs (panels 3A and 3B, respectively; gTRCA eigenvalue = 2.21 and 2.66, both significantly reproducible at the group level, p < 0.002). These components exhibited a sequence of positive and negative deflections at different latencies (PM: 13, 39, 81, and 120 ms; M1: 13, 55, 123, 172, 262, and 354 ms). Both stimulations involved lateralized channels located more frontally for PM and more posteriorly for M1. On the other hand, Figure 3C and 3D shows the distributions of individual GMFPs. Most subjects exhibited significant responses lasting up to 308 ms for PM and 285 ms for M1, as indicated by the median GMFPs (blue traces). Figure 3E highlights timepoints where significant evoked responses differ between PM and M1 TEPs in each subject. In at least 50% of subjects, significant amplitude differences persisted for up to 262 ms, whereas only a small fraction of participants showed no differences between targets (orange area). These differences primarily involved frontoparietal EEG channels over the stimulated hemisphere (right side of Figure 3E). Similar results were observed for the phase analysis (Figure 3F).

**Figure 3.**
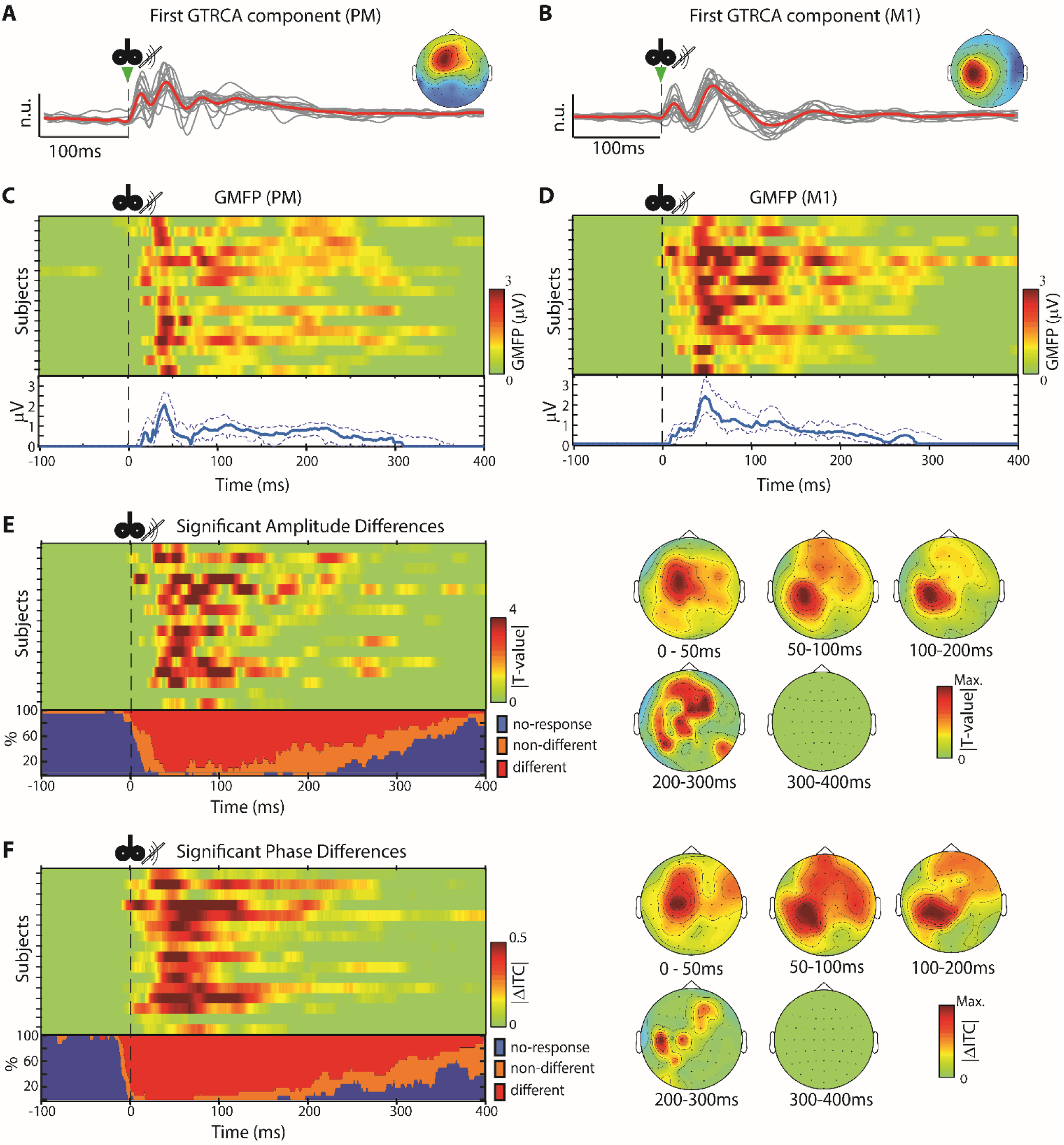
Significant differences in amplitude and phase between premotor and motor TEPs are confirmed both at the group and individual levels. **(A, B).** Waveforms of the most reproducible component at the group level, as extracted by gTRCA, for both premotor (A) and motor (B) TEPs, along with their average scalp maps representing the strength and direction of the component at each EEG channel. Gray lines are the principal gTRCA components of individual subjects, and the red line is the average component across subjects. **(C, D)** GMFP of significant TEPs evoked by premotor (C) and motor (D) stimulation for each subject (n=16). Solid blue traces represent median GMFP values, with dashed lines indicating interquartile ranges. **(E, F)** Results of the cluster-based statistical tests for each subject. Heatmaps show the time courses of average significant amplitude (E) and phase (F) differences between EEG responses to premotor and motor stimulation. Each line represents the average absolute T-values (E) and ITC differences (F) across significant clusters for one subject. The bottom plots illustrate, for each time sample, the percentage of subjects presenting evoked responses, in amplitude (E) and phase (F), that are significantly different between premotor and motor stimulation (red), evoked responses that are not significantly different between targets (orange), and the percentage of subjects with no evoked response (blue). Topographical distribution of absolute T-values (E) and ITC differences (F) for significant clusters (median across subjects), averaged over different time windows (0-50 ms, 50-100 ms, 100-200 ms, 200-300 ms, 300-400 ms), are shown on the right.

Overall, these findings show that eTMS to nearby cortical targets can evoke EEG responses characterized by spatiotemporal evolutions that differ up to about 300 ms at both group and single-subject levels.

### RS-TMS evokes stereotypical EEG responses

We sought counterfactual evidence that TMS-related sensory co-stimulations can indeed result in EEG responses that, unlike TEPs, do not significantly differ over both space and time. To this aim, we first compared EEG responses evoked by RS-TMS of the same two scalp targets during eTMS using the same analyses as for TEPs. As such, Figure 4A and 4B show the grand average EEG responses to RS-TMS applied to PM and M1, respectively. RS-TMS evoked significant activity between 47-358 ms for PM and 48-342 ms for M1. Visual inspection of scalp topographies relative to GMFP peaks revealed similar activations, primarily over central electrodes, for both targets. This was supported by cluster-based statistical analysis, which revealed no significant differences in either amplitude or phase (Figure 4C and 4D). Additionally, the most reproducible gTRCA components were qualitatively similar across conditions, exhibited low group-level reproducibility (gTRCA eigenvalues: 0.81 for PM and 0.85 for M1, both p < 0.002), and were characterized by slow, stereotyped waveforms peaking over central electrodes (Figure 4E and 4F). Notably, RS-TMS applied over PM and M1 failed to elicit significant responses in 3 and 6 subjects, respectively. In the remaining participants, RS-TMS evoked low-amplitude responses (Figure 4G and 4H, and Figure S4B). Crucially, the late responses (100–400 ms) to RS-TMS were significantly smaller than the corresponding TEPs (LMM, β = −0.22, t(82) = −3.527, p < 0.001), as reflected by the GMFP analysis (Figure S5 and Table S6).

**Figure 4.**
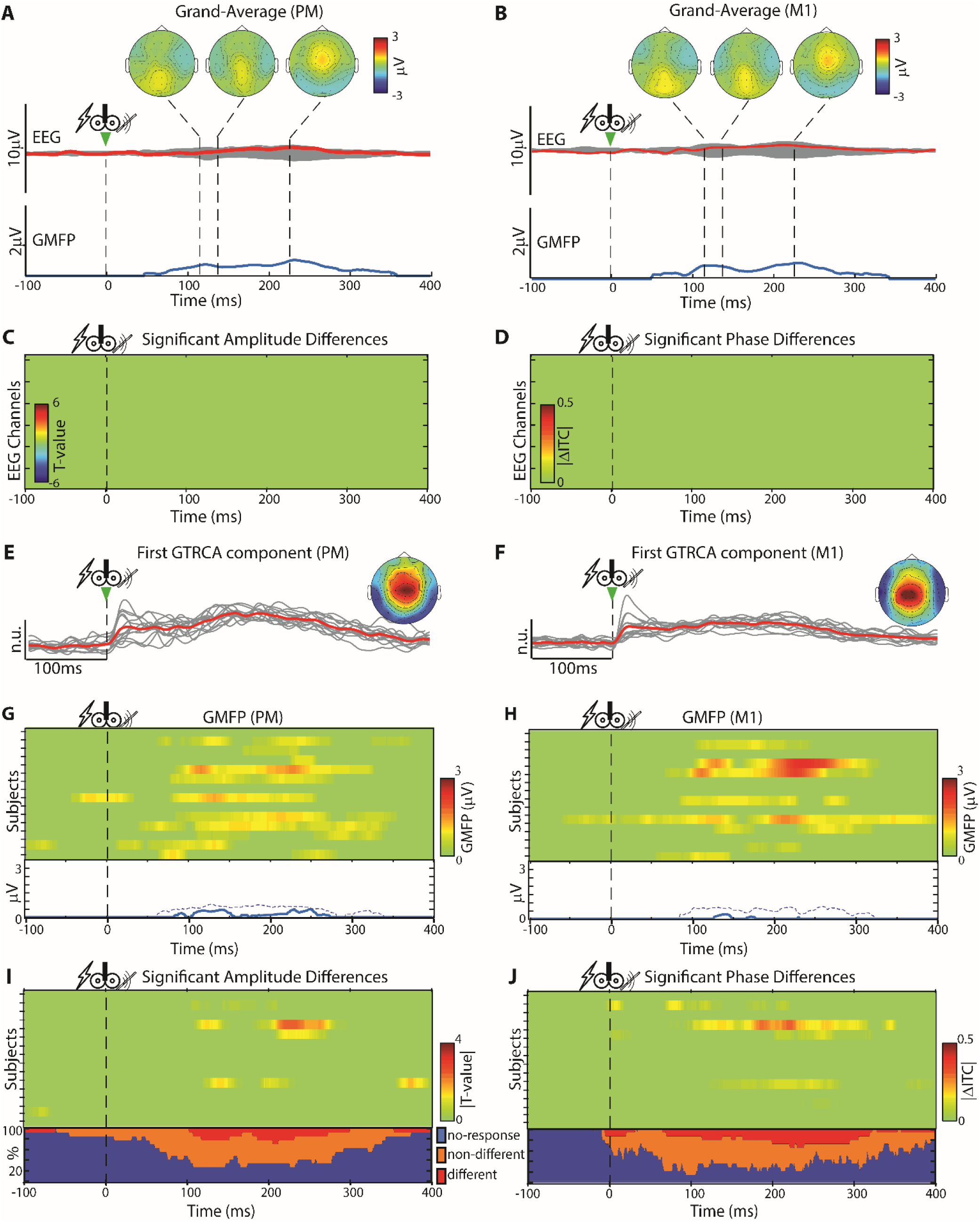
Realistic sham TMS evokes stereotypical EEG responses. This figure replicates the analyses from Figures 2 and 3 using EEG responses evoked by RS-TMS. **(A, B)** Grand-average evoked responses following RS-TMS for premotor (A) and motor (B) stimulation, including butterfly plots, highlighted channels, GMFP traces, and topographical maps at GMFP peaks, as in Figures 2A and 2B. **(C, D)** There are no significantly different clusters in either amplitude (C) or phase (D) when comparing grand-average EEG responses evoked by premotor and motor RS-TMS. **(E, F)** Waveforms of the principal gTRCA component, along with their average scalp maps, as in Figures 3A and 3B. (G, H) GMFP of significant evoked potentials for each subject (n=14), including median and interquartile GMFP values, as in Figures 3C and 3D. **(I, J)** Results of the cluster-based statistical tests for each subject, as in Figures 3E and 3F. Throughout all time points, only a minority of subjects (maximum 5 out of 14) show responses to RS-TMS that are significantly different between premotor and motor stimulation.

We further compared EEG responses to RS-TMS applied to PM and M1 using cluster-based analysis at the single-subject level. Only 5 of 14 subjects showed significant differences in amplitude or phase domains (Figure 4I and 4J). In contrast to TEPs, over 70% of subjects showed no differences in spatiotemporal patterns between targets, as indicated by the sum of the blue (no response) and orange areas (responses not differing significantly between targets).

Overall, we show that, in the absence of direct cortical engagement, sensory stimulation evokes EEG responses whose spatiotemporal evolution is stereotypical, involves central electrodes and does not significantly differ between stimulation targets, both at the group and single-subject levels.

### HI-ES evokes large but stereotypical EEG responses

Previous studies have shown that RS-TMS can result in evoked potentials of significantly lower amplitude compared to TEPs [29,40]. Thus, to maximize EEG responses to electrical scalp stimulation we performed HI-ES over both stimulation targets and compared the ensuing EEG responses using the same analyses as for TEPs and RS-TMS. In this case, average responses to HI-ES emerged at late latencies (Figure 5A and 5B). PM stimulation evoked significant responses between 43 and 490 ms, while responses to M1 stimulation occurred between 22 and 444 ms. Scalp topographies relative to GMFP peaks showed contralateral activations around 70 ms followed by central activations for both targets. Unlike RS-TMS, the cluster-based test identified significant amplitude differences between targets, but, unlike TEPs, these differences occur in a narrow time window (150–210 ms) and primarily over central electrodes (Figure 5C). Comparisons in the phase domain revealed significant clusters from 112 to 209 ms and 299 to 368 ms (Figure 5C and 5D), overlapping with the same scalp regions as the amplitude domain. The most reproducible gTRCA component evoked by HI-ES was similar across targets (Figure 5E and 5F; gTRCA eigenvalue = 1.73 and 1.62 for PM and M1, respectively, both with p < 0.002), peaking over central electrodes, resembling those obtained with RS-TMS.

**Figure 5.**
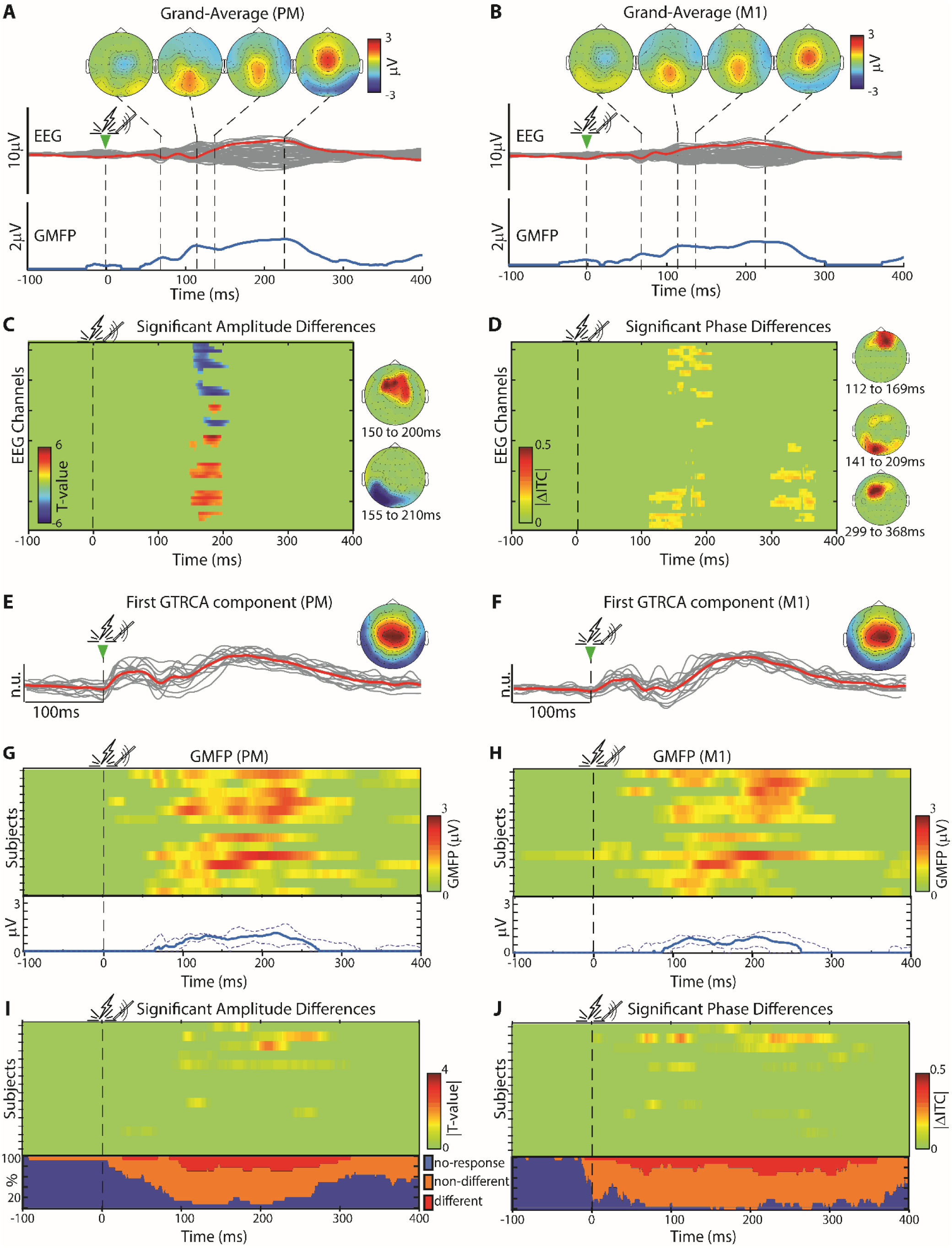
High-intensity electrical stimulation elicits only marginal amplitude and phase differences between premotor and motor stimulation. This figure shows the results of the same analyses from the previous figures using the responses evoked by HI-ES. **(A, B)** Grand-average evoked potentials for premotor (A) and motor (B) stimulation, including butterfly plots, highlighted channels, GMFP traces, and topographical maps at GMFP peaks, as in Figures 2A and 2B. **(C, D)** Two significantly different clusters were found in amplitude (C) and three in phase (D) when comparing grand-average EEG responses evoked by premotor and motor HI-ES. These panels show significant clusters’ heatmaps and topographical distributions as in Figures 2C and 2D. **(E, F)** Waveforms of the principal gTRCA component evoked by HI-ES, along with their average scalp maps, as in Figures 3A and 3B. **(G, H)** GMFP of significant potentials evoked by HI-ES for each subject (n=14), including median and interquartile GMFP values, as in Figures 3C and 3D. **(I, J)** Results of the cluster-based statistical tests for each subject, as in Figures 3E and 3F. Similarly to RS-TMS (Figure 4), consistent differences between premotor and motor stimulation are observed in only a minority of subjects (a maximum of 5 out of 14; red area panels I and J).

Analysis of GMFP at the individual level confirmed that HI-ES evoked larger responses than RS-TMS at late latencies (Figure 5G and 5H, and Figure S4C), with amplitudes resembling those of TEPs (LMM, β = −0.01, t(82) = −0.161, p = 0.87) (Figure S5 and Table S6). However, larger HI-ES responses did not result in differences between PM and M1. Specifically, the percentage of subjects with a significant response to HI-ES (orange area of Figure 5I and 5J) was similar to that observed with TMS (Figure 3I and 3J), while the percentage showing significant differences between targets (red area of Figure 5I and 5J) was comparable to that observed with RS-TMS (Figure 4I and 4J).

This third set of results shows that EEG responses to electrical scalp stimulation, even when maximized, involve central electrodes and do not significantly differ between stimulation targets at both the group and single-subject levels.

### The hemispheric asymmetry of late evoked responses to eTMS, but not to RS-TMS nor HI-ES, discriminates between stimulation targets

Results so far show that EEG responses evoked by eTMS of different, yet nearby, cortical targets exhibit distinct voltage waveforms and scalp topographies, even at late latencies. On the contrary, EEG responses to RS-TMS and HI-ES at late latencies are stereotyped and nonspecific, involving mainly central electrodes, regardless of the stimulation target. Results also suggest that lateralization of late responses is a key feature for differentiating TEPs from PM and M1 targets, but not EEG responses by RS-TMS or HI-ES. To systematically test this, we computed a simple index to quantify EEG response lateralization (Supplementary Methods 11) at early (0–100 ms) and late (100–400 ms) latencies. Figure 6 shows the performance of such index quantified by Receiver Operating Characteristic (ROC) analysis. TEPs reported an area under the curve (AUC) of 0.69, increasing to 0.91 for the early and the late time windows, respectively. On the contrary, the lateralization of EEG responses to RS-TMS and HI-ES cannot distinguish between targets in both time windows (AUC for 0–100 ms: 0.51 for RS-TMS and 0.56 for HI-ES; AUC for 100–400 ms: 0.57 for RS-TMS and 0.51 for HI-ES).

**Figure 6.**
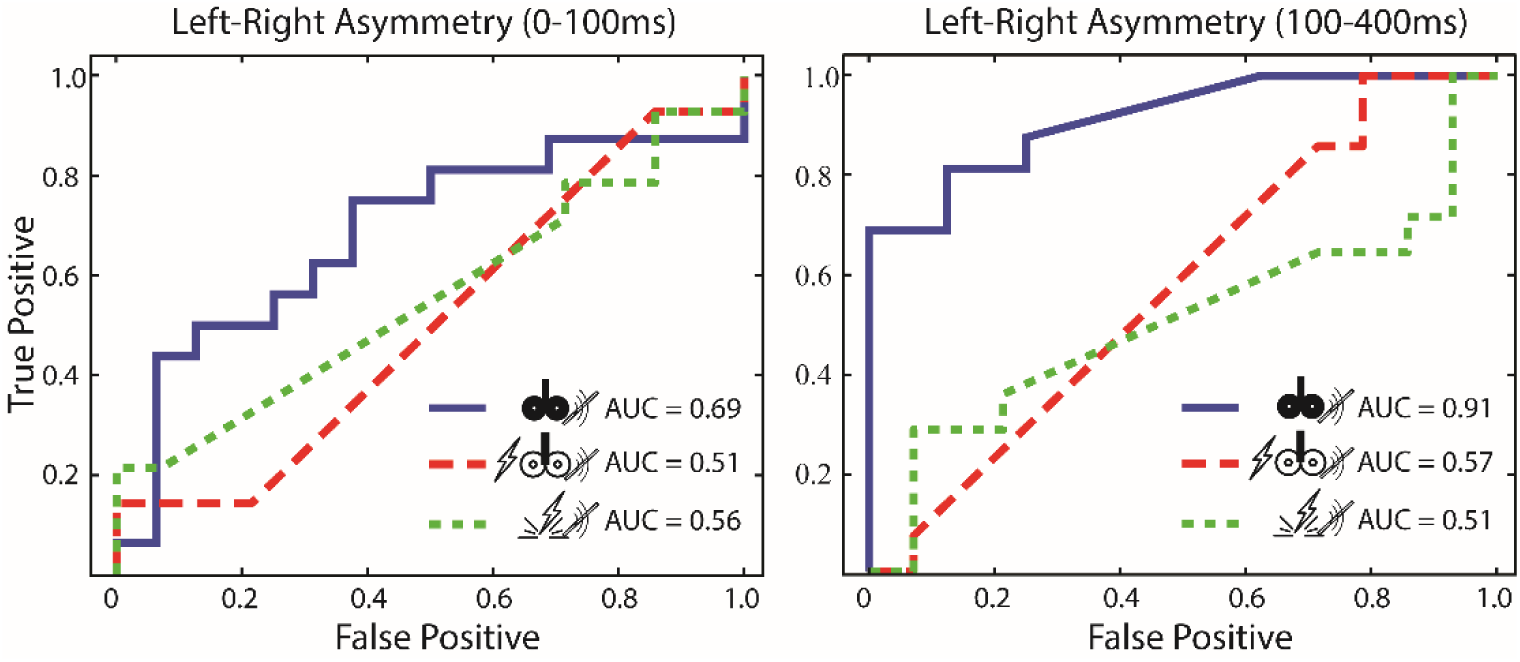
Hemispheric asymmetry of late evoked responses discriminates between motor and premotor stimulation in TEPs but not in peripheral evoked potentials. For each recording session, significant differences in the mean field power between left and right EEG channels were used to compute an asymmetry index representing the lateralization of the evoked response (Supplementary Methods 11). This figure displays the results of the receiver operating characteristic (ROC) analysis for discriminating between premotor and motor evoked responses using the asymmetry index averaged across early (0-100 ms) and late latencies (from 100 to 400 ms). ROC curves are shown for eTMS (solid blue line), RS-TMS (dashed red line), and HI-ES (dashed green line), along with the corresponding values of the area under the curve (AUC).

## DISCUSSION

The dominant view in TMS-EEG research is that only early TEP components reflect genuine cortical responses to TMS, while later components mainly arise from sensory confounds [19–22,24–28,40,41]. Here, we tested whether the spatiotemporal evolution of TEPs, including their later components, can reflect a genuine cortical activation and hence provide specific information about the targeted cortical circuits. To address this question, we compared TEPs from two adjacent cortical targets in a group of healthy participants, challenging the null hypothesis that their spatiotemporal features do not differ. This hypothesis was specifically tested under conditions designed to optimize the direct cortical effects of TMS (eTMS) while minimizing sensory co-stimulation. As a counterfactual comparison, we applied the same analytical approach to EEG responses elicited by realistic sham TMS (RS-TMS) and by high-intensity electrical scalp stimulation (HI-ES), both lacking direct cortical activation.

We found that eTMS of one premotor and one motor cortical target results in significantly different responses throughout a 300 ms time window and across large scalp regions. These differences are particularly evident over the stimulated hemisphere and detectable at the single-subject level (Figures 2, 3, and S4). Conversely, both RS-TMS and HI-ES elicit EEG responses that are either indistinguishable between targets or show differences limited in both time and space (Figures 4 and 5, and S4). These findings contrast with previous reports and suggest that sensory responses do not primarily drive TEPs, even at late latencies. Several factors may explain this discrepancy.

Previous studies suggest that TEP components can be significantly affected by TMS-related AEPs [29,42], especially at late latencies. When noise masking is absent or ineffective, late TEPs are visibly influenced by the N100–P180 complex [7,17,43], typically prominent over central EEG electrodes and potentially reducing site-specificity [10]. Consistent with this, we observed that TMS discharge in sham position produces AEPs with similar spatiotemporal features across stimulation sites (Figures S3A and S3B). As previously shown [29,44,45], we confirm that AEPs can be abolished by optimizing noise-masking parameters [11,34] with ad hoc procedures (Supplementary Methods 2), in both RS-TMS and eTMS, at the group and, crucially, the single-subject level (Figures S3C and S3D).

In addition to TMS-related AEPs, the lack of control over TMS effectiveness in engaging cortical targets may explain discrepancies in the literature [16]. Here, we used a real-time TEP visualization tool [11] to ensure that the initial (< 50 ms) cortical response peaked in EEG channels beneath the TMS coil with a preset amplitude > 6 µV after averaging 20 epochs. These early, high-amplitude local TEPs represent a prerequisite (i.e., the independent variable) to effectively activate the target neuronal population [46–49] and engage circuit-specific interactions [50,51](i.e., the dependent variable of interest).

By adhering to this methodological criterion, we found that the spatiotemporal evolution of TEPs differed significantly between the two cortical targets, with amplitude and phase differences extending up to 300 ms at the single-subject level. This is particularly noteworthy given the proximity of the targets and their shared role in the motor circuit [52,53]. The robustness of these differences is further supported by a novel analytical approach, gTRCA, which identifies the most reproducible TEP components across epochs and subjects for a given set of stimulation parameters [38,39]. These components showed marked differences in waveform and scalp topography between targets (Figure 3), consistent with prior findings of site-dependent variations in waveforms, frequency content, and cortical activation [30–32,35,54–59]. Such differences likely reflect the recruitment of distinct circuits, as shown by recent studies combining TMS with intracranial recordings [60] and cortical source reconstruction integrated with connectivity modelling [61]. Specifically, primary motor cortex stimulation produced slow, lateralized activations (Figure S4A), consistent with previous studies using subthreshold intensities [29]. These late components are unlikely due to sensory input and may arise from cortico-thalamic loops, as suggested by their absence in thalamectomized patients [62]. They can also be modulated by coil orientation [63] and motor tasks [64], reflecting their dependence on the engagement and the state of the target cortical circuits, respectively. Conversely, stimulation of medial premotor cortex evoked late (∼200 ms), slow components more prominent at central electrodes. A recent multi-scale study of premotor TMS responses suggests these late components reflect engagement of specific medial cortical circuits [65]. In particular, thalamic rebound activity makes a major contribution to the ∼200 ms midline TEPs elicited by premotor TMS, whose amplitude is strongly modulated by motor tasks that alter the state of thalamocortical neurons. These EEG findings are supported by analogous observations in human LFP recordings, as well as in rodent LFP and unit data [65] and similar late components also appear with intracortical electrical stimulation—which lacks sensory co-stimulation—further underscoring their origin in cortico-thalamo-cortical circuits [66]

After demonstrating significant, long-lasting differences in premotor and motor TEPs across time and space, we examined counterfactual evidence and found (when present) stereotyped spatiotemporal activations for RS-TMS and HI-ES, both delivered without direct cortical activation. Late TEP components are often attributed to somatosensory-evoked potentials [67], arising from craniofacial muscle contractions or activation of scalp somatosensory receptors and nerve branches [68]. To mimic the somatosensory input from TMS discharge, RS-TMS combined scalp electrical stimulation (matched to TMS-related perception) and TMS in sham position preserving vibratory components [43]. Under this condition, EEG responses were dominated by a central N100-P200 complex, with no significant differences between premotor and motor targets. Notably, nearly half the participants showed no significant EEG response to RS-TMS (Figure 4), consistent with prior sham-TMS studies [29,40]. This likely reflects the limited density of cutaneous somatosensory afferents in the scalp [69], which constraints the ability of peripheral stimulation to generate detectable cortical EEG responses. Also relevant are TMS-EEG studies in brain-injured patients—with preserved somatosensory pathways—who showed absent or minimal EEG responses when sensory confounds were minimized and TMS targeted structurally lesioned cortex [70,71].

We then raised the intensity of electrical scalp stimulation to the maximum tolerable level below pain perception, eliciting significant EEG responses for both premotor and motor targets in all but one participant. Despite substantial intensity differences, EEG responses to HI-ES showed stereotypical components similar to those from RS-TMS (Figures 4, 5, and S4), and crucially, were invariant across stimulation targets (Figure 6). The average waveform presented a ∼70 ms component with a topography slightly contralateral to the stimulation site, consistent with sensory responses to forehead electrical stimulation [72]. This was followed by two later components resembling the N100-P200 complex, typical of multimodal vertex responses to salient sensory inputs [73]. These components were slightly larger for the premotor target, and only in a minority of subjects (Figure 5), possibly reflecting greater cortical responsiveness to scalp regions near the face [74]. Overall, these findings suggest that, unlike eTMS, high-intensity somatosensory stimulation of different scalp areas activates the same neural network for detecting and processing salient stimuli.

Lastly, it has been suggested that TMS-related somatosensory activations may involve scalp or muscle nociceptors [68]. In our study, the scalp muscle contractions were controlled by coil position adjustments [11]. Accordingly, only one out of 16 participants perceived pain during eTMS over the motor cortex (Figure S1), which rules out significant contributions of pain-related evoked potentials to late TEP components.

While the contribution of somatosensory responses to late TEP components cannot be entirely excluded, our findings suggest such influence is likely negligible. This interpretation is reinforced by ROC analysis, which showed that response lateralization following eTMS—but not RS-TMS or HI-ES—reliably distinguished stimulation targets (Figure 6), with the strongest discriminative power in the 100–400 ms window. These results were obtained using rigorous procedures to minimize artifacts and ensure effective cortical engagement—an approach that not only reduces TMS-related artifacts during acquisition but also lessens the need for extensive preprocessing to extract TEPs [75,76](Figure S2). Notably, this strategy may not generalize to all TMS-EEG studies, such as those targeting lateral (e.g., dorsolateral prefrontal cortex) or posterior (e.g., cerebellum) structures. In such cases, spurious activations must be carefully evaluated and addressed.

Overall, our results indicate that, when TMS effectiveness is maximized and peripheral confounds are adequately controlled, the technique evokes distinct and long-lasting EEG responses that reflect genuine activation of targeted cortical regions. These results challenge previous assumptions and highlight the utility of TMS-EEG as a robust tool for engaging intrinsic local circuit dynamics and investigating large-scale functional interactions within corticothalamic networks.

## Acknowledgments

The authors are thankful to Michele Colombo and Renzo Comolatti for their insightful discussions regarding data analysis. We also thank Alessandra Dalla Vecchia and Giorgio Mariotti for their assistance during data acquisition.

## Author Contributions

Conceptualization: M.F., S.R., A.G.C., M.R.

Data curation: M.F., A.G.C.

Formal analysis: M.F., B.A.C., A.G.C.

Funding acquisition: M.F., A.P., S.S., S.C., M.M., M.R.

Investigation: M.F., S.R., G.F., G.H., S.D.A., M.S., A.V., S.P., M.R.

Methodology: M.F., S.R., A.G.C., M.R.

Software: M.F., S.R., B.A.C., A.G.C.

Supervision: M.F., S.C., S.S., M.M., A.G.C., M.R.

Writing – original draft: M.F., S.R., A.G.C., M.R.

Writing – review and editing: M.F., S.R., B.A.C., E.M., A.P., S.P., S.S., S.C., M.M., A.G.C., M.R.

## Data Availability Statement

The data that support the findings of this study as well as individual and group-level data are available upon request.

