## Supplementary Material for "The spatiotemporal evolution of TMS-evoked potentials reflects direct cortical activation"

**SUPPLEMENTARY METHODS**

1. Neuronavigated Transcranial Magnetic Stimulation ... 3
2. Noise Masking Configuration, AEPs, and auditory sham controls ... 3
3. Optimization of TMS parameters for maximization of cortical activation and minimization of confounds ... 3
4. Electrical stimulation parameters ... 4
5. MEP Monitoring ... 4
6. Numerical Rating Scale (NRS) ... 4
7. EEG preprocessing ... 4
8. Independent Component Analysis ... 5
9. Cluster analysis ... 5
10. Statistics to assess significant evoked potentials ... 5
11. Indexing the degree of lateralization of evoked potentials ... 6

**SUPPLEMENTARY FIGURES**

Figure S1: Psychometric report of the perceptual features as reported by the study participants after each measurement ... 7
Figure S2: Effect of preprocessing on TEPs and peripheral evoked potentials ... 8
Figure S3: Responses evoked by TMS click with and without noise masking ... 9
Figure S4: Premotor and motor evoked responses in a representative subject ... 11
Figure S5: Amplitude of late components evoked by eTMS, RS-TMS and HI-ES ... 13

**SUPPLEMENTARY TABLES**

Table S1: Demographic characteristics ... 14
Table S2: Numerical Rating Scale (NRS) ... 15
Table S3: Stimulation Intensities ... 16
Table S4: Linear mixed-effects model predicting Discomfort Ratings ... 17
Table S5: Linear mixed-effects model predicting Pain Ratings ... 18
Table S6: Linear mixed-effects model predicting the amplitude of late evoked responses ... 19
**SUPPLEMENTARY REFERENCES ... 20**

### SUPPLEMENTARY METHODS

#### 1. Neuronavigated Transcranial Magnetic Stimulation

Cortical areas were targeted using an air-cooled, neuronavigated figure-of-eight TMS coil (mean and outer winding diameters: 50 mm and 70 mm; stimulation area: 0.68 cm²; NBS 5.2.5, Nexstim, Finland), delivering biphasic pulses of 280 µs duration. The neuronavigation system allowed the visual identification of cortical targets using T1-weighted individual MRI scans obtained at either 1.5 T or 3 T (Philips and Siemens Prisma, respectively). It also allowed for the estimation of the TMS-induced electric field intensity in V/m (E-field) and the localization of its maximum (E-field hot spot) over the cortical surface [1]. Stimulation parameters such as the coil position and orientation were stored in the neuronavigation system and subsequently used for locating the electrodes for electrical stimulation. Most importantly, the neuronavigation system allowed for the maintenance of constant stimulation parameters throughout the recording session (ensuring control of the hot spot location and pausing the stimulation when off-target) and for the digitization of the EEG electrodes' position, which ensured that the relative position between the EEG cap and subject's head did not differ across TMS and sham recording sessions [2]. Additionally, it can display TMS Motor Evoked Potentials (MEP).

#### 2. Noise Masking Configuration, AEPs, and auditory sham controls

The TMS coil was tilted at 90° in the sagittal plane, directly over the head vertex, and in contact with the scalp ("sham" position). Ten to twenty TMS pulses were delivered at 80% of the maximum output to familiarize the subject with the TMS coil "click." Then, the subject was asked to wear inserted earplugs and was asked to report every time he/she heard a TMS "click" while a noise masking generated by a custom-made tool [3] was playing. Every time the subject correctly identified the TMS click, the noise parameters (resampling frequency and balance between click-based noise and white noise, sound volume) were optimized. The procedure was repeated iteratively, adjusting the parameters of the noise masking until the subject no longer reported hearing the TMS "click."

To verify that AEPs were effectively abolished by the noise masking, at the end of each TMS recording session we recorded AEPs with and without noise masking during TMS "sham" stimulation of both cortical targets at the intensity used for TEPs, and as described in [3]. For each condition, at least 100 pulses were delivered with the coil in the "sham" position over the targeted cortical area. EEG preprocessing followed the pipeline described for TMS-EEG data (see section 7). Good epochs were 122 ± 36 (premotor) and 138 ± 29 (motor) for sham TMS without noise masking and 120 ± 31 (premotor) and 134 ± 31 (motor) for sham TMS with noise masking (values are reported as mean ± standard deviation). No more than 2 channels were rejected across all subjects.

#### 3. Optimization of TMS parameters for maximization of cortical activation and minimization of confounds

Cortical targets were first visually selected based on the subject's cortical anatomy [4], as displayed by the NBS system. We positioned the TMS coil over the subject's scalp to ensure the E-field hot spot was located over the crown of the selected gyrus (superior frontal gyrus for PM and precentral gyrus around the "hand knob" for M1). Then, one TMS pulse was delivered at a stimulation intensity set around 120 V/m to check for the activation of TMS-induced cranio-facial muscles. In case of scalp muscle twitching, the orientation of the coil, the intensity of the stimulation, and eventually the selected cortical spot was recursively slightly adjusted. To verify the effectiveness of the stimulation parameters in activating the cortex, we delivered 10 to 20 TMS pulses and visually checked that early (< 50 ms) TEP components were largest in the EEG leads underneath the TMS coil using the rt-TEP tool [5]. If these components displayed a peak-to-peak amplitude below the desired threshold (set at 6 µV in this study), we adjusted the stimulation parameters (increase in stimulation intensity, or slight coil rotation and/or translation). We also avoided the activation of the cortico-spinal tract by ensuring that MEPs were not elicited (see Section 5). Finally, when present, decay artifacts were minimized by fixing the impedances or by rearranging the electrode's lead wire [6] before starting a recording session. Electro-Gel was used for TMS-EEG recordings (Electro-Cap International Inc.)[7] to ensure the recording stability.

#### 4. Electrical stimulation parameters

Electrical stimulation of the scalp, delivered as square-wave pulses lasting 200 μs with a maximum compliance voltage set at 200 V, was administered using a DS7A electrical stimulator (Digitimer Ltd., Welwyn Garden City, UK) via bipolar Ag/AgCl cup electrodes (10 mm diameter, Spes Medica S.r.l.). Stimulating electrodes were inserted through additional holes made in the EEG cap fabric and positioned about 2.5 cm apart along an axis approximating the location where the TMS coil made contact with the scalp and the direction of the TMS-induced current [8,9] as stored in the neuronavigation system. To minimize the electrical stimulation artifact and prevent electrical bridging, stimulating electrodes were positioned to avoid contact with the recording electrodes and were secured in place using Ten20 conductive paste (Weaver & Company, Aurora, Colorado, USA). To further minimize the amplitude and duration of the electrical decay artifact, the polarity of the stimulating electrodes was inverted with every pulse (alternating polarity mode) throughout the recording, a procedure similar to [8]. Moreover, Abralyt HiCl gel (EASYCAP GmbH) was used to reduce high-amplitude, long-lasting decay artifacts induced by the electrical stimulation. Indeed, the high chloride concentration in Abralyt HiCl gel ensured stability of DC recordings with Ag/AgCl electrodes [10]. The intensity of the electrical stimulation was set as follows. For the RS-TMS, the subject was stimulated with increasing intensities until reporting that the intensity was comparable to the TMS intensity. To ensure accurate comparisons between the perceived intensity of the current electrical stimulation with the original TMS, electrical pulses were alternated with TMS pulses of the original intensity used in the subject. For HI-ES, the current intensity was set by gradually increasing the delivered current to the maximum tolerable level without generating pain.

#### 5. MEP Monitoring

To prevent TMS-evoked motor contractions, we continuously monitored MEPs using six electrodes electromyogram (EMG) placed over the right hand muscles (APB, the FDI, and the FLB muscles recorded through Ag-AgCl self-adhesive electrodes placed in a belly-tendon). We used a 6-channel eXimia electromyography (EMG) system with a sampling rate of 3000 Hz and a cutoff of 500 Hz for low-pass filtering [11]. If MEPs were present, the coil was slightly translated and/or rotated until they disappeared, and/or related epochs were discarded during pre-processing.

#### 6. Numerical Rating Scale (NRS)

After each recording session, the subject was asked to give a score on a numerical rating scale (NRS) ranging from 0 to 10 to report the perceived loudness of the coil's click sound (from "no clicks heard" to "I heard them all"), the perceived discomfort and the pain (from "no perception" to "maximal perception") that the stimulation was producing. At the end of each RS-TMS and HI-ES recording session, in addition to the described NRS, the participant was asked whether the perceived stimulus was qualitatively perceived as similar to the real TMS. Here we report questions that were asked: 1) Did you perceive the intensity of the Sham stimulation as similar to the TMS stimulation? If not, which one did you perceive as more intense? 2) Was the tactile perception of the electrical stimulation, as a whole, different from that of TMS? If yes, how so?

#### 7. EEG preprocessing

EEG data were analyzed in MATLAB R2016b. Bad channels and epochs with discontinuities were discarded by visual inspection. TMS pulse and discharge artifacts were removed by replacing the interval around the stimulus (-2 to 8 ms) with the preceding interval (-12 to -2 ms) and applying a moving-average filter (5th order) between 6 and 10 ms. DC fluctuations were removed with a 0.5 Hz high-pass filter for RS-TMS and HI-ES and with a 1 Hz high-pass filter for TMS-EEG (3rd order Butterworth). The choice of applying different high-pass filters to the eTMS and the RS-TMS and HI-ES data is due to the presence of the decay artifact produced by the electrical stimulation and the need to minimally distort it to allow the Independent Component

Analysis (ICA) algorithm for better identification. Data were epoched from -1000 to 1000 ms around the simulations. Good epochs were 207 ± 39 for eTMS, 298 ± 47 for RS-TMS, and 299 ± 43 for HI-ES (values are reported as mean ± standard deviation), with no more than 3 channels rejected. To remove artifacts from eye movements, muscle activity, and decay, ICA was performed on data referenced to the average of the signal recorded at all EEG electrodes (section 8 below). If present, mixed ICA components were considered good. Finally, data were low-pass filtered at 45 Hz (3rd order Butterworth), high-pass filtered at 1 Hz (3rd order Butterworth) in the case of RS-TMS and HI-ES, downsampled to 1 kHz, and segmented from -600 to 600 ms around the stimulus to remove possible edge effects. To avoid bias in subsequent statistical comparisons, the first 135 consecutive good epochs were used for constructing the final evoked potentials in each recorded session (this number corresponds to the minimum number of good epochs across all experimental conditions, targets, and participants).

#### 8. Independent Component Analysis

Independent Component Analysis (ICA) is a blind source separation algorithm that can extract independent components from multidimensional data. However, as reported by Metsomaa and colleagues [12], time-locked multi-trial non-stationary EEG data may not fulfill the assumption of statistical independence. In this vein, reducing TMS-evoked artifacts during data acquisition, such as evoked scalp muscles and decays, not only allows for the correct use of the ICA but also reduces the impact of the analysis pipeline on the final TEP. In this work, the ICA procedure was applied to remove spontaneous eye movements, saccades, and spontaneous muscle activity.

On the other hand, the decay artifacts generated by the electrical stimulation in sham conditions are unavoidable. These artifacts may differ from epoch to epoch due to a degrading electrode conductance or, as in this case, due to the polarity reversion performed at each epoch. These differences among epochs are detected by the ICA and isolated in a maximum of two components.

#### 9. Cluster analysis

Evoked responses between premotor and motor stimulations were compared using a nonparametric permutation test, corrected for multiple comparisons through a two-step cluster-based statistics (open-source FieldTrip Toolbox [13]). First, we tested the effect at each spatiotemporal sample, establishing a threshold to identify samples as cluster members. T-statistics were used for amplitude comparisons, while phase differences were estimated using the absolute value of the difference in normalized analytical signals extracted from the broadband Hilbert transform [14]. Significant spatiotemporal samples ($p < 0.05$; two-sided parametric test for voltages and one-sided, nonparametric test for phases) were grouped based on temporal and spatial adjacency (with a minimum of two EEG channels per cluster). The sum of first-level statistics within each cluster served as cluster-level statistics and was compared to the maximum distribution of values obtained after randomizing data across conditions (using Monte Carlo approximation with 1000 random permutations). Clusters were considered significant when the observed summed statistics exceeded 95% of the values resulting from random permutations.

#### 10. Statistics to assess significant evoked potentials

To assess the statistical significance of EEG responses in amplitude, we used a surrogate data approach based on circularly shifting trial sequences [15]. For each EEG session (whether individual subject or grand-average), we generated 1000 surrogate sessions by randomly shifting the temporal series of the original trials, while preserving the temporal sequence of the real voltages in surrogate EEG data. The shifting factor was drawn from a uniform distribution, and all EEG channels within a trial were shifted by the same amount. We then compared the original evoked signal to this set of surrogate data using the “clusterstat” method of the FieldTrip toolbox in the same voltage-based nonparametric clustering analysis described above. The resulting binary matrices, representing spatiotemporal samples with significant evoked responses, were used to compute the global mean field power (GMFP), the topographical distributions of evoked responses, and to filter the results of the voltage-based cluster analysis comparing premotor and motor stimulation.

Significant phase locking across trials was estimated by calculating the broadband phase locking factor (PLF)[16]. The PLF was computed for each EEG session by averaging the normalized analytic signal, obtained from the Hilbert transform, across trials. For each electrode, the mean pre-stimulus phase locking was used to construct a Rayleigh cumulative distribution, representing the null hypothesis of no phase locking across trials. PLF values were considered significant when exceeding 99% of the values in the Rayleigh distribution (Bonferroni corrected). The resulting binary matrices were used to filter the results of the phase-based cluster analysis described previously.

#### 11. Indexing the degree of lateralization of evoked potentials

To assess the topographical asymmetry of the evoked potentials, we compared EEG responses between hemispheres for each recording session. EEG electrodes were grouped into frontal, central, and parietal regions of interest (ROIs) per hemisphere (Frontal left: F1, F3, FC1, FC3; Frontal right: F2, F4, FC2, FC4; Central left: C1, C3, CP1, CP3; Central right: C2, C4, CP2, CP4; Parietal left: P1, P3, PO3; Parietal right: P2, P4, PO4). Mean field power (MFP) differences at these ROIs were calculated and compared to a distribution of maximum differences values obtained from 1000 random permutations of trials across hemispheres. Significant differences were classified as Left > Right (L > R) or Right > Left (R > L) and averaged across the three ROIs and over early (from 0 to 100ms) and late latencies (from 100 to 400 ms). An asymmetry index was calculated for each time window by subtracting total L > R power from total R > L power, indicating higher evoked power in the left or right hemisphere, respectively. For each experimental condition (TMS, RS-TMS and HI-ES), this index was used to discriminate between stimulation targets using the area under the curve (AUC) of the Receiver Operating Characteristic (ROC) analysis.

### SUPPLEMENTARY FIGURES

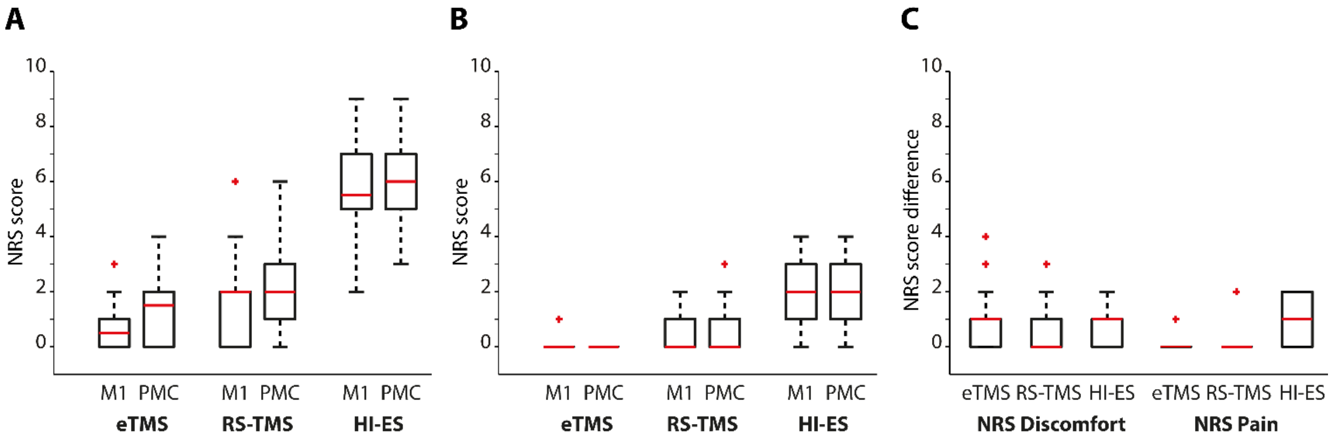


**Figure S1: Psychometric report of the perceptual features as reported by the study participants after each measurement.**

**(A,B)** NRS values of discomfort (A) and pain (B) as reported by the study participants after eTMS, RS-TMS, and HI-ES measurements over premotor and motor areas (0 = no discomfort/pain; 10 = maximum discomfort/pain). In the case of perceived discomfort, a linear mixed-effects model (LMM) analysis revealed a significant main effect of stimulation type, but no significant effect of stimulation area ($\beta = 0.65$, $t(82) = 1.37$, $p = 0.17$, Table S4). With respect to eTMS, discomfort NRS values were on average 1.03 points higher during RS-TMS ($t(82) = 2.17$, $p = 0.03$) and 4.7 points higher during HI-ES ($t(82) = 9.99$, $p = 2x10^{-16}$). Similar results were obtained for pain, with no significant effect of stimulation area ($\beta = -0.0625$, $t(82) = -0.227$,$p = 0.82$, Table S5) but a significant effect of stimulation type. Pain NRS values were on average 0.36 points higher during RS-TMS ($t(82) = 1.27$, $p = 0.2$) and 2.1 points higher during HI-ES ($t(82) = 7.245$, $p = 2x10^{-11}$) when compared to eTMS. For both outcomes, there were no significant interactions between area and type of stimulation ($p > 0.6$). **(C)** Paired differences between premotor and motor NRS are shown on the right for each stimulation type and the two NRS, indicating how much the stimulation was perceived to be similar between targeted spots. Each boxplot displays the median (red line), and the first and third quartiles (bottom and top edges of the box). Outliers are indicated by the red crosses, and whiskers extend from the bound of the box to the data points not considered outliers.

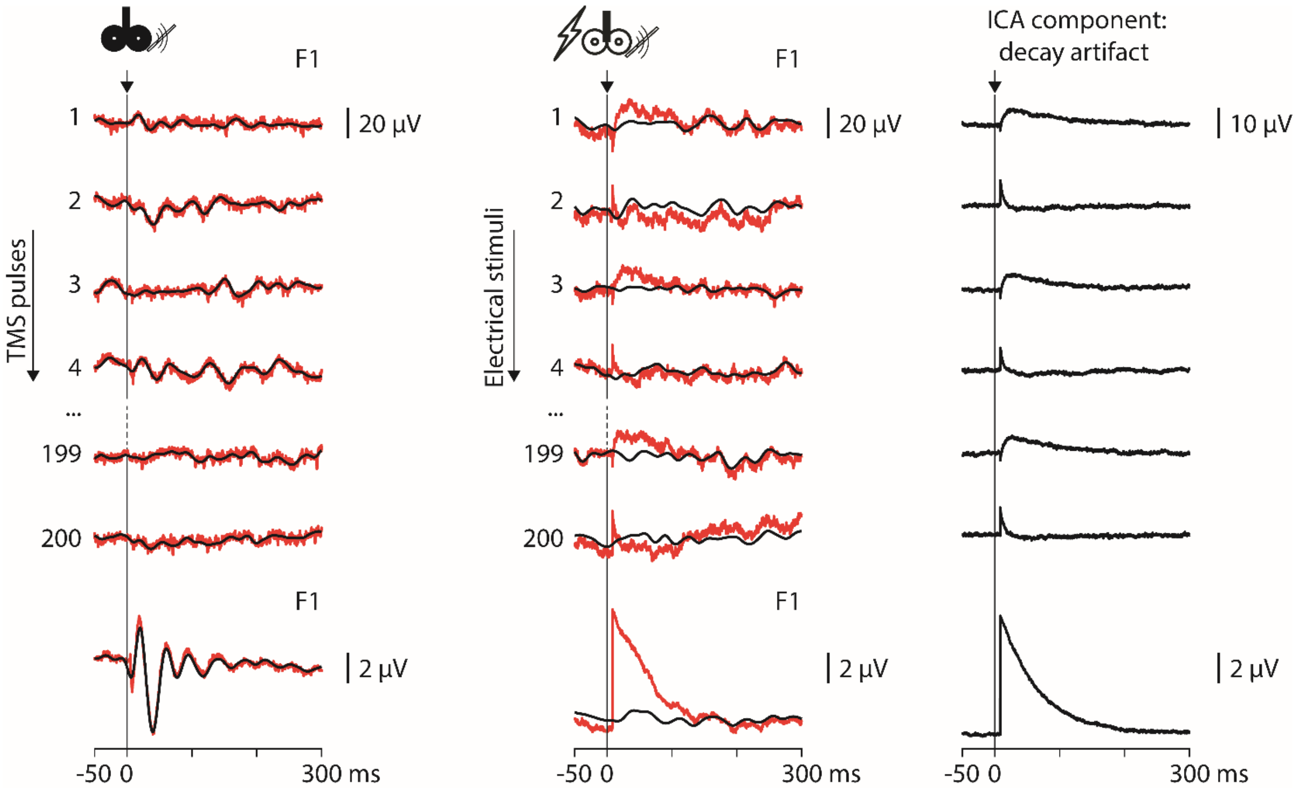


**Figure S2: Effect of preprocessing on TEPs and peripheral evoked potentials**

The EEG response evoked on the F1 channel by eTMS of the premotor area before (red traces) and after ICA components removal and 45 Hz low-pass filtering (black traces) are shown on the left. The upper rows show brain activity following eTMS at each epoch after artifact removal, high-pass filtering at 1 Hz, and re-referencing to the average, while the bottom row shows the average evoked response. Similar to the left panel, epochs and average electrical scalp evoked response are shown in the middle panel. The difference between red and black traces highlights the decay artifact introduced by electrical stimulation and its effective removal through the pre-processing pipeline. The EEG signal reconstructed from the ICA component identified as a decay artifact is shown on the right panel. The alternation between positive and negative waveforms observable in each epoch reflects the polarity reversion of the electrical stimulation.

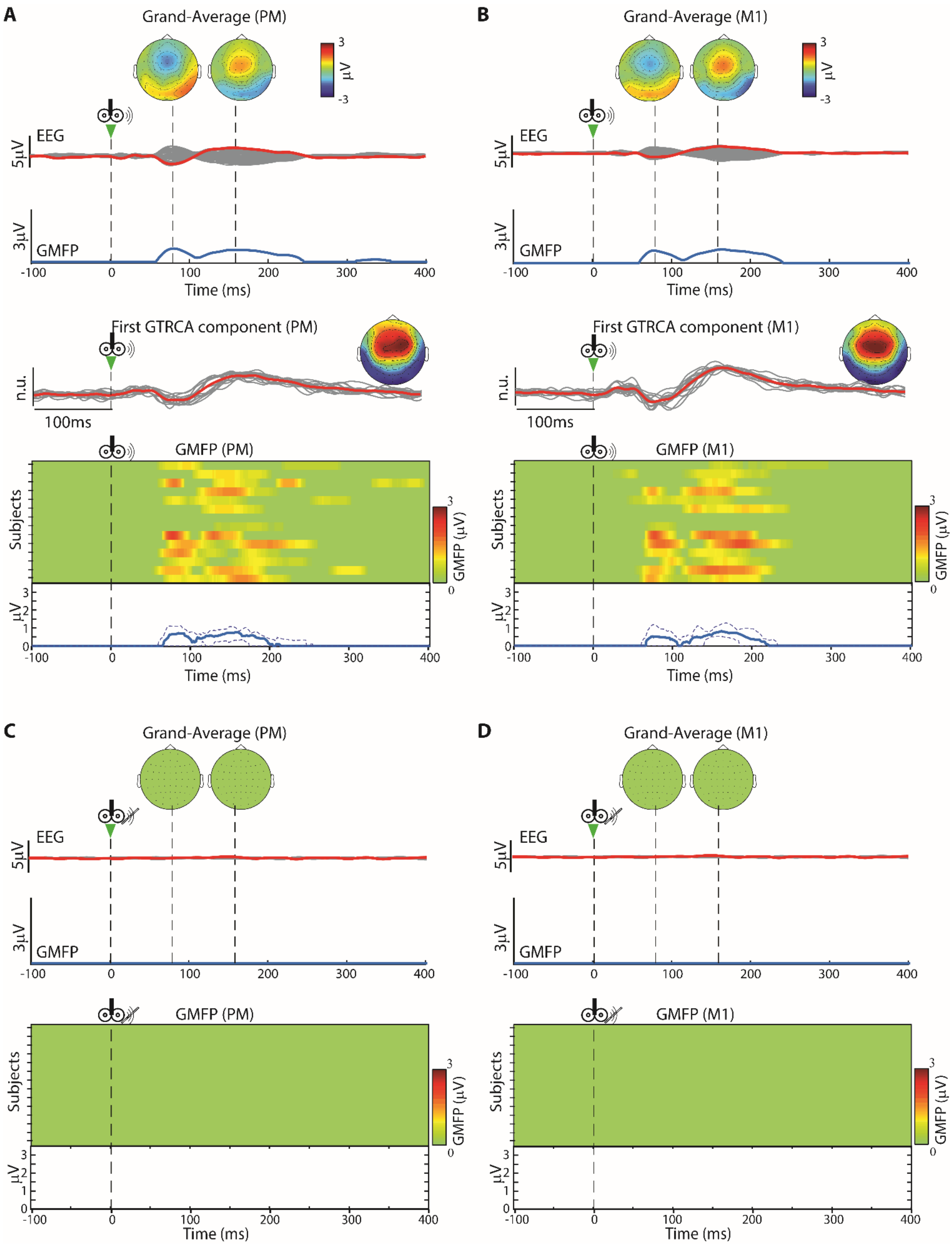


**Figure S3: Responses evoked by TMS click with and without noise masking.**

TMS clicks without noise masking consistently evoke significant auditory EEG responses. In contrast, TMS clicks with noise masking do not evoke significant auditory EEG responses**. (A, B)** From top to bottom: Grand average auditory response evoked by the TMS click while stimulating above the premotor (A) and motor (B) cortex with the coil in the sham position, along with topographic voltage distribution at peak times (heads above) and the corresponding significant GMFP over time (trace below); significant gTRCA component; significant GMFP over time for each subject (heatmap), along with median (blue solid line) and interquartile range (blue dashed line) across subjects. (**C, D)** Grand average EEG response evoked by the TMS click above the premotor (C) and motor (D) cortex with noise masking. Bottom row: significant GMFP over time for each subject. There were no significant gTRCA components at the group level, and no significant evoked response was observed in any subject.

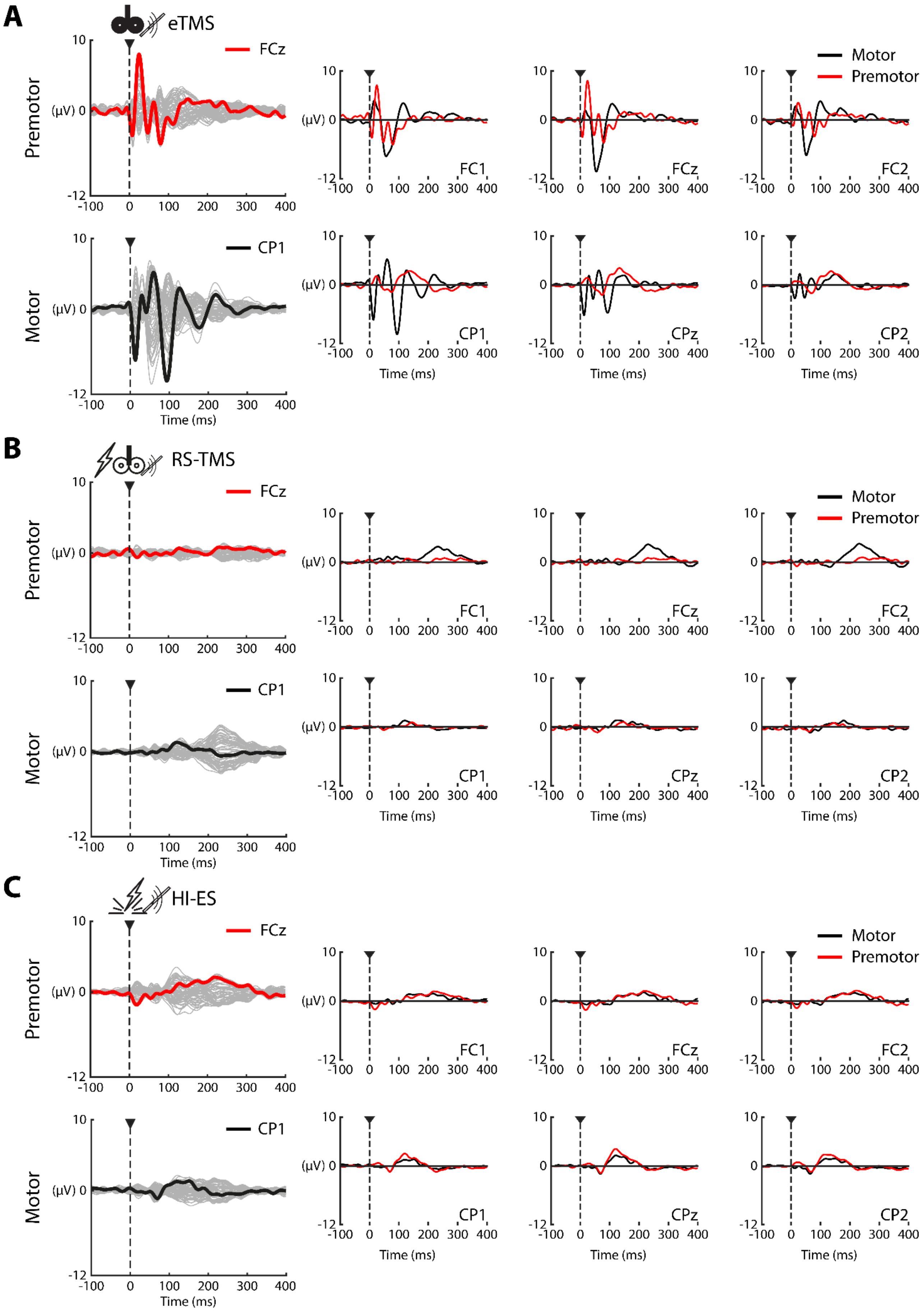
A
eTMS
B
RS-TMS
C
HI-ES
Premotor
Motor
FCz
CP1
Motor
Premotor
FC1
FCz
FC2
CP1
CPz
CP2
(μV) 0
10
-12
-100
0
100
200
300
400
Time (ms)

**Figure S4: Premotor and motor evoked responses in a representative subject**

EEG responses evoked by eTMS (A), RS-TMS (B), and HI-ES (C) of premotor (top) and motor cortex (bottom) across EEG channels (gray) are shown on the left. The channel with the largest response evoked by the eTMS (premotor: FCz, red; motor: CP1, black) is shown in each condition. Responses of premotor (red) and motor cortex (black) in the 6 frontocentral and central channels close to the midline are superimposed on the right. The responses evoked by eTMS of the premotor and motor cortex exhibit different peaks and waveforms bigger closer to the stimulation site. RS-TMS evoked a slow response while stimulating M1 which is bigger in the frontocentral channels, while the response evoked by stimulating the PM appears small and not different from the pre-stimulus activity. HI-ES evoked a slow, stereotyped response different from pre-stimulus activity that peaked around central scalp EEG electrodes.

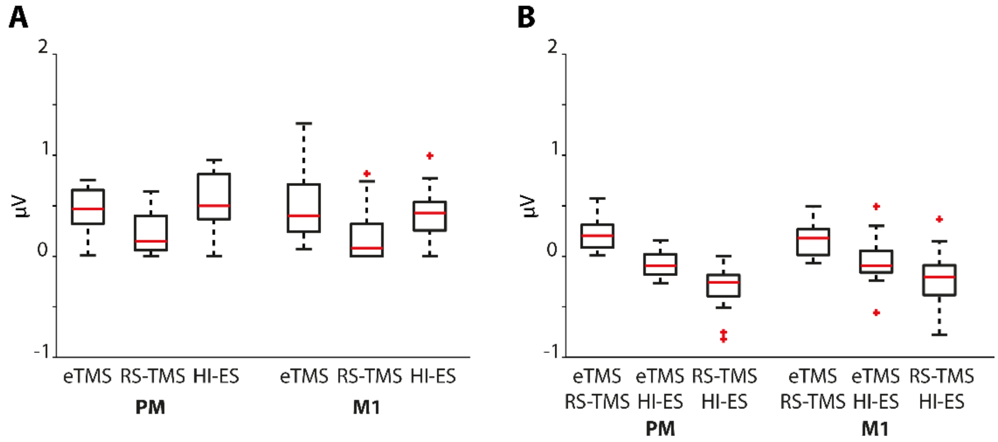


**Figure S5: Amplitude of late components evoked by eTMS, RS-TMS and HI-ES**

**(A)** Amplitudes of late components, calculated as the average GMFP between 100 and 400 ms post-stimulation, are displayed for each stimulation type grouped by targeted area. (**B**) Paired amplitude differences between stimulation types. With respect to eTMS, RS-TMS evoked significantly smaller late responses ($\beta = -0.22$, $t(82) = -3.527$, $p < 0.001$, Table S6), while HI-ES evoked similar responses ($\beta = -0.01$, $t(82) = -0.161$, $p = 0.87$). There were no significant main effects of stimulation area ($\beta = -0.025$, $t(82) = -0.412$, $p = 0.68$) and no significant interaction between area and type of stimulation ($p > 0.17$). Boxplots display median (red line), first and third quartiles (bottom and top edges of the box), and outliers (red crosses). The whiskers extend to the data points that are not considered outliers.

### SUPPLEMENTARY TABLES

| Subject ID | Age | Gender | Handedness |
|---|---|---|---|
| **S01** | 28 | F | R |
| **S02** | 37 | M | R |
| **S03** | 40 | M | L |
| **S04** | 32 | F | L |
| **S05** | 35 | M | R |
| **S06** | 36 | M | R |
| **S07** | 27 | M | R |
| **S08** | 51 | M | R |
| **S09** | 45 | M | R |
| **S10** | 28 | F | R |
| **S11** | 25 | M | R |
| **S12** | 35 | M | R |
| **S13** | 36 | F | L |
| **S14** | 27 | M | R |
| **S15** | 41 | M | R |
| **S16** | 46 | M | R |

**Table S1: Demographic characteristics.**

For each subject (rows) the ID, age, gender, and handedness (columns) are reported.

| | NRS Discomfort | | | | | | NRS Pain | | | | | |
|---|---|---|---|---|---|---|---|---|---|---|---|---|
| | eTMS | | RS-TMS | | HI-ES | | eTMS | | RS-TMS | | HI-ES | |
| **Subject ID** | **PM** | **M1** | **PM** | **M1** | **PM** | **M1** | **PM** | **M1** | **PM** | **M1** | **PM** | **M1** |
| **S01** | 1 | 3 | 4 | 2 | 7 | 5 | 0 | 0 | 0 | 0 | 2 | 1 |
| **S02** | 0 | 0 | 1 | 1 | 8 | 8 | 0 | 0 | 0 | 0 | 4 | 4 |
| **S03** | 2 | 1 | 2 | 0 | 7 | 6 | 0 | 0 | 0 | 0 | 3 | 1 |
| **S04** | 0 | 1 | 2 | 2 | 3 | 3 | 0 | 1 | 0 | 2 | 1 | 3 |
| **S05** | 0 | 0 | 2 | 2 | 6 | 5 | 0 | 0 | 2 | 2 | 4 | 3 |
| **S06** | 2 | 1 | 1 | 0 | 4 | 2 | 0 | 0 | 0 | 0 | 0 | 0 |
| **S07** | 0 | 0 | 2 | 2 | 6 | 7 | 0 | 0 | 0 | 0 | 2 | 3 |
| **S08** | 4 | 0 | 4 | 4 | 6 | 7 | 0 | 0 | 2 | 0 | 4 | 2 |
| **S09** | 0 | 0 | 1 | 0 | 3 | 3 | 0 | 0 | 0 | 0 | 2 | 4 |
| **S10** | 4 | 3 | n.a | n.a | n.a | n.a | n.a | n.a | n.a | n.a | n.a | n.a |
| **S11** | 2 | 1 | 2 | 2 | 6 | 5 | 0 | 0 | 0 | 0 | 1 | 2 |
| **S12** | 1 | 0 | n.a | n.a | n.a | n.a | n.a | n.a | n.a | n.a | n.a | n.a |
| **S13** | 0 | 0 | 6 | 6 | 9 | 9 | 0 | 0 | 3 | 1 | 3 | 2 |
| **S14** | 3 | 0 | 3 | 4 | 5 | 6 | 0 | 0 | 1 | 1 | 3 | 3 |
| **S15** | 2 | 1 | 3 | 0 | 6 | 6 | 0 | 0 | 0 | 0 | 2 | 2 |
| **S16** | 2 | 2 | 0 | 0 | 5 | 5 | 0 | 0 | 0 | 0 | 0 | 0 |

**Table S2: Numerical Rating Scale (NRS)**

Individual NRS scores rating the perceived discomfort (left) and pain (right) for each stimulation type (eTMS, RS-TMS, and HI-ES) and targeted site (PM - premotor and M1 - motor). NRS spans from 0 (“no perception”) to 10 (“maximal perception”).

| | eTMS (%MO) | | RS-TMS (mA) | | HI-ES (mA) | |
|---|---|---|---|---|---|---|
| **Subject ID** | **PM** | **M1** | **PM** | **M1** | **PM** | **M1** |
| **S01** | 57% | 58% | 6,45 | 4,24 | 11,00 | 6,80 |
| **S02** | 50% | 44% | 8,05 | 3,20 | 11,30 | 5,20 |
| **S03** | 45% | 46% | 3,10 | 2,15 | 4,90 | 4,00 |
| **S04** | 50% | 31% | 2,50 | 4,81 | 6,00 | 7,30 |
| **S05** | 53% | 42% | 5,02 | 5,96 | 9,50 | 9,00 |
| **S06** | 40% | 32% | 4,50 | 3,10 | 5,60 | 4,50 |
| **S07** | 42% | 43% | 5,83 | 5,13 | 6,20 | 5,90 |
| **S08** | 48% | 50% | 3,18 | 3,17 | 6,15 | 6,99 |
| **S09** | 54% | 51% | 8,00 | 7,00 | 18,00 | 20,00 |
| **S10** | 45% | 34% | n.a | n.a | n.a | n.a |
| **S11** | 60% | 63% | 6,80 | 5,35 | 7,20 | 7,60 |
| **S12** | 42% | 40% | n.a | n.a | n.a | n.a |
| **S13** | 45% | 52% | 5,68 | 3,80 | 8,40 | 4,30 |
| **S14** | 58% | 59% | 5,25 | 5,46 | 9,50 | 12,50 |
| **S15** | 49% | 57% | 4,24 | 4,70 | 12,10 | 8,18 |
| **S16** | 44% | 31% | 5,80 | 4,91 | 18,00 | 14,00 |

**Table S3: Stimulation Intensities**

The stimulation intensities are reported for each subject (rows), stimulation type, and site (columns). eTMS intensities are reported as the percentage of the stimulator's maximal output (%MO), while RS-TMS and HI-ES intensities are reported as mA.

| Fixed effects | Estimate (β) | SE | 95% CI | t(82) | p |
|---|---|---|---|---|---|
| Intercept (Target = M1, Condition = eTMS) | 0.813 | 0.389 | [0.040, 1.585] | 2.09 | 0.040 |
| Target = PM | 0.625 | 0.456 | [-0.281, 1.531] | 1.37 | 0.174 |
| Condition = RS-TMS | 1.033 | 0.475 | [0.088, 1.978] | 2.17 | 0.033 |
| Condition = HI-ES | 4.747 | 0.475 | [3.802, 5.692] | 9.99 | <0.001 |
| Interaction (PM, RS-TMS) | -0.054 | 0.667 | [-1.380, 1.273] | -0.08 | 0.936 |
| Interaction (PM, HI-ES) | -0.339 | 0.667 | [-1.666, 0.987] | -0.51 | 0.612 |

| Random effects | Estimate (SD) | 95% CI |
|---|---|---|
| Intercept (Participant) | 0.869 | [0.520, 1.452] |
| Residual | 1.288 | [1.093, 1.518] |

**Table S4: Linear mixed-effects model predicting Discomfort Ratings**

Results of Linear Mixed Effects model for numerical rate scale (NRS) of discomfort. Results reported for each fixed effect are the unstandardised coefficient (β̂), its standard error (SE), 95 % confidence interval (CI), *t*-statistic (t) and p-value (p). Positive coefficients indicate higher discomfort relative to the reference condition (M1 under eTMS). Random-effect and residual standard deviations (SD) quantify between-participant variability and within-participant error, respectively. Model specification: *NRS (discomfort) ~ Target × Condition + (1 | Participant)*. Maximum-likelihood estimation was used (N = 88 observations from 16 participants); denominator degrees of freedom were 82. Model fit statistics: AIC = 330.07, BIC = 349.89, log-likelihood = -157.03, deviance = 314.07.

| Fixed effects | Estimate (β) | SE | 95% CI | t(82) | p |
|---|---|---|---|---|---|
| Intercept (Target = M1, Condition = eTMS) | 0.0625 | 0.2188 | [-0.373, 0.487] | 0.2856 | 0.7759 |
| Target = PM | -0.0625 | 0.2756 | [-0.611, 0.486] | -0.227 | 0.821 |
| Condition = RS-TMS | 0.364 | 0.287 | [-0.206, 0.935] | 1.270 | 0.207 |
| Condition = HI-ES | 2.079 | 0.287 | [1.508, 2.649] | 7.245 | <0.001 |
| Interaction (PM, RS-TMS) | 0.205 | 0.404 | [-0.597, 1.008] | -0.509 | 0.612 |
| Interaction (PM, HI-ES) | 0.134 | 0.404 | [-0.669, 0.937] | -0.332 | 0.741 |

| Random effects | Estimate (SD) | 95% CI |
|---|---|---|
| Intercept (Participant) | 0.398 | [0.222, 0.713] |
| Residual | 0.779 | [0.663, 0.916] |

**Table S5: Linear mixed-effects model predicting Pain Ratings**

Results reported for each fixed effect are the unstandardised coefficient (β̂), its standard error (SE), 95 % confidence interval (CI), *t*-statistic (t) and p-value (p). Positive coefficients indicate higher pain relative to the reference condition (M1 under eTMS). Random-effect and residual standard deviations (SD) quantify between-participant variability and within-participant error, respectively. Model specification: *NRS (pain) ~ Target × Condition + (1 | Participant)*. Maximum-likelihood estimation was used (N = 88 observations from 16 participants); denominator degrees of freedom were 82. Model fit statistics: AIC = 235.94, BIC = 255.76, log-likelihood = -109.97, deviance = 219.94.

| Fixed effects | Estimate (β) | SE | 95% CI | t(82) | p |
|---|---|---|---|---|---|
| Intercept (Target = M1, Condition = eTMS) | 0.481 | 0.0669 | [0.348, 0.614] | 7.194 | <0.001 |
| Target = PM | -0.0246 | 0.0596 | [-0.143, 0.094] | -0.412 | 0.681 |
| Condition = RS-TMS | -0.220 | 0.0625 | [-0.345, -0.096] | -3.527 | <0.001 |
| Condition = HI-ES | -0.010 | 0.0625 | [-0.134, 0.114] | -0.161 | 0.872 |
| Interaction (PM, RS-TMS) | 0.0158 | 0.0873 | [-0.158, 0.189] | 0.181 | 0.857 |
| Interaction (PM, HI-ES) | 0.121 | 0.0873 | [-0.0529, 0.294] | 1.384 | 0.170 |

| Random effects | Estimate (SD) | 95% CI |
|---|---|---|
| Intercept (Participant) | 0.208 | [0.139, 0.311] |
| Residual | 0.169 | [0.143, 0.199] |

**Table S6: Linear mixed-effects model predicting the amplitude of late evoked responses.**

Results reported for each fixed effect are the unstandardised coefficient (β̂), its standard error (SE), 95 % confidence interval (CI), *t*-statistic (t) and p-value (p). Positive coefficients indicate higher GMFP of late (100-400ms) evoked responses relative to the reference condition (M1 under eTMS). Random-effect and residual standard deviations (SD) quantify between-participant variability and within-participant error, respectively. Model specification: *GMFP ~ Target × Condition + (1 | Participant)*. Maximum-likelihood estimation was used (N = 88 observations from 16 participants); denominator degrees of freedom were 82. Model fit statistics: AIC = -12.384, BIC = 7.434, log-likelihood = 14.192, deviance = -28.384.
